# TransBind2: Improving Transcription Factor-DNA Binding Prediction with Multimodal Data and Bidirectional Cross Attention

**DOI:** 10.64898/2026.09.07.749913

**Authors:** Shreya Basnet, Jianlin Cheng

**Affiliations:** Department of Electrical Engineering & Computer Science, University of Missouri, Columbia, MO 65211, USA; NextGen Precision Health, University of Missouri, Columbia, MO 65211, USA

## Abstract

Accurate genome-wide prediction of transcription factor (TF)–DNA binding remains challenging because many models focus mainly on DNA sequence and overlook chromatin context and TF structure. We previously developed TransBind, a protein-aware model that combines TF and DNA representations through cross-attention. Here, we introduce TransBind2, which improves on TransBind in several ways. It incorporates DNase-seq accessibility and genome mappability tracks as additional input, uses a biomodal protein language model (ProstT5) to capture both TF sequence and structure, and applies bidirectional cross-attention so DNA and protein features can refine each other. We also frame prediction as binary classification of individual *<*DNA bin, TF, cell type*>* triplets, allowing the model to generalize to new TFs and cell types. Across 690 human ChIP-seq experiments covering 161 TFs and 91 cell types, TransBind2 achieves a macro AUROC of 0.9648 and AUPR of 0.4215, outperforming TransBind and other baselines, with a ≥12.67% relative AUPR gain. The model trained on human data also performs well in cross-species zero-shot prediction on mouse data. Saliency analysis shows that it can identify TF-binding peaks with a median error of 12–38 base pairs (bps) despite being trained on window-level labels. Ablation studies further show that TF structure, chromatin accessibility, and bidirectional attention each improve performance. Overall, these results show that combining TF structure with chromatin context leads to more accurate and generalizable TF–DNA binding predictions.

## Introduction

Transcription factors (TFs) regulate gene expression by binding short, sequence-specific elements in the genome, thereby establishing the transcriptional programs that govern cellular identity, development, and function [14]. Chromatin immunoprecipitation followed by sequencing (ChIP-seq) remains the gold standard for mapping TF–DNA interactions genome-wide [5, 27], but its cost, material requirements, and dependence on high-quality antibodies mean that experimentally derived binding maps cover only a small fraction of the vast combinatorial TF-DNA interaction space defined by TFs, cell types, and regulatory contexts. Computational prediction of TF–DNA binding has therefore become an essential complement to experimental profiling.

Early computational approaches, including position weight matrices, hidden Markov models, and biophysical binding-site discovery methods [10, 19, 20], captured core sequence preferences but could not model longer-range context or higher-order regulatory grammar. Deep learning subsequently transformed the field. Early convolutional architectures such as DeepBind learned motif-like features directly from DNA sequence [2], while multitask models such as DeepSEA and hybrid CNN–recurrent architectures such as DanQ improved the modelling of regulatory sequence context and longer-range dependencies [23, 37]. Later work incorporated attention mechanisms to improve interpretability [6, 22]. Complementary studies also demonstrated that DNA structural and biophysical properties, including minor groove width, roll, helix twist, and DNA breathing dynamics, contribute to binding specificity [3, 7, 35, 38]. Beyond DNA shape descriptors, structure-based methods have also been applied directly to TF–DNA complexes. A recent study combined AlphaFold 3-predicted TF–DNA structures [1] with the physics-based scoring tool FoldX [8] to assess the allele-specific effects of noncoding variants on TF–DNA binding. This structural modelling approach reproduced experimentally observed binding preferences without requiring TF-specific training. However, its predictive accuracy remained lower than that of sequence-based models, and performance varied substantially across different TFs [12]. Collectively, these studies illustrate a broader shift from purely sequence-based representations toward incorporating the physical and structural determinants of TF–DNA recognition. In parallel, genome foundation models have emerged as another major direction for TF–DNA binding prediction. Models such as DNABERT-2 have learned transferable sequence representations from large genomic corpora [39], and hybrid approaches such as EPBD×DNABERT-2 combine such representations with physical DNA binding features to further improve prediction [17].

Despite these advances, most TF–DNA binding predictors remain fundamentally DNA-centric. However, TF–DNA recognition is determined jointly by the properties of both the DNA and the TF, including DNA sequence context as well as TF sequence, structure, and biophysical characteristics [28]. Reflecting this shift toward protein-aware modelling, BTFBS [15] took an initial step by representing TFs explicitly during prediction. However, it relied on small curated binding-site databases and encoded TFs only as raw amino-acid tokens, without leveraging contextual protein embeddings or structural information [15]. Protein language models pretrained on large sequence corpora, such as ESM-2 and its DNA-binding-protein-adapted variant ESM–DBP, offer a more expressive alternative for representing TFs [26, 36]. Building on this idea, we previously introduced TransBind, a protein-aware architecture that integrates ESM–DBP-derived TF embeddings with a convolutional/BiLSTM/ transformer-based [32] DNA encoder through cross-attention, allowing each TF embedding to selectively attend to genomic regions according to its binding properties [4]. TransBind was evaluated on 690 ChIP-seq experiments spanning 161 TFs and 91 human cell types. It outperformed existing DNA-centric methods, recovered 160 known binding motifs in JASPAR [25] directly from its convolutional filters, and supported label-zero-shot prediction for TFs excluded from training.

Contemporaneously, TFBindFormer introduced a hybrid cross-attention transformer that likewise conditions DNA representations on TF-specific embeddings derived from protein sequence and structure-derived tokens, further underscoring the move toward protein-aware, structurally informed binding models [18]. Like TransBind, however, TFBindFormer relies on DNA sequence and protein features alone, without considering chromatin accessibility or genome uniqueness (or sequence mappability information), both of which are known to influence in vivo TF occupancy [11, 29].

In addition, TransBind’s TF representation is reduced to a single vector obtained by mean-pooling embeddings from a sequence-only protein language model, thereby discarding residue-level information and any explicit representation of protein structure, despite the inherently structural nature of TF–DNA recognition [21, 33]. Furthermore, its cross-attention mechanism is unidirectional: DNA features attend to protein features, but not vice versa, limiting the ability of the two modalities to iteratively refine one another. Finally, TransBind is formulated as a multilabel classifier over a fixed set of 690 TF–cell type experiments. As a result, generalization to unseen TFs requires a separately designed zero-shot variant rather than arising naturally from modelling pairwise TF–DNA compatibility.

Here we present TransBind2, which addresses each of these limitations. First, we augment the DNA sequence input with position-aligned DNase-seq accessibility signal and genome uniqueness (mappability) scores. FactorNet established the use of DNase-seq accessibility as a position-aligned input channel alongside DNA sequence [24]. DeepGRN later extended this framework by incorporating a Duke-style genome uniqueness (mappability) track and demonstrated through feature ablation that DNase-seq accessibility was the single most informative predictor of TF–DNA binding [6]. We adopt the same accessibility-plus-uniqueness combination [9, 11] here, computed at the resolution of full 1000 bp genomic bins, to provide TransBind2 with direct information about regulatory context and sequence repetitiveness. Second, we replace the sequence-only, averaged ESM–DBP embedding of TF used by TransBind with the embedding generated by ProstT5 [34]. ProstT5 jointly encodes amino-acid sequence together with AlphaFold-predicted structural information by learning correspondences between protein sequence and Foldseek’s 3Di structural alphabet, thereby producing structure-aware TF representations. [13, 16, 31]. Third, we replace TransBind’s unidirectional attention with a symmetric, bidirectional cross-attention mechanism in which DNA and protein representations iteratively refine each other before prediction. Fourth, we reformulate the learning problem as binary pairwise classification over individual *<*DNA bin, TF, cell type*>* triplets, so that generalization to novel TFs and cell types is a native property of the model rather than a separately trained variant.

The results show that TransBind2 substantially outperforms previous methods across the large majority of TFs evaluated. It also generalizes in a cross-species zero-shot setting to mouse ChIP-seq data despite being trained exclusively on human data and localizes TF-binding peaks with a median error of 12–38 bp using input×gradient saliency maps despite being trained only with window-level binding labels. Together, these results indicate that explicitly modeling chromatin context and TF structure, combined with bidirectional cross-modal attention, yields a more accurate and more generalizable framework for genome-wide TF–DNA binding prediction.

## 1 MATERIALS AND METHODS

### 1.1 Multimodal Data

#### 1.1.1 DNA data

TransBind2 uses the same genomic sequence bins as TransBind, derived from the EPBDXDNA dataset [17] and the GRCh37 (hg19) human reference genome. The chromosome-based data split is also retained, with chromosomes 8 and 9 reserved for testing, chromosome 7 used for validation, and the remaining chromosomes used for training. All genomic regions are represented as 1,000 bp bins (windows). The primary difference between TransBind2 and TransBind lies in the formulation of the prediction task. In TransBind [4], each genomic bin is treated as a single training example associated with a 690-dimensional multilabel vector, where each dimension corresponds to a specific transcription factor (TF)–cell type experiment. The shortcoming of this multilabel formation is that the number of predicted classes is tied to the number of unique TFs in the training data, making it difficult to generalize to new TFs unseen in the training data. In contrast, TransBind2 reformulates the problem as binary classification, such that each example consists of a single *<*DNA bin, TF, cell type*>* triplet with a binary label indicating whether the TF binds to that genomic region in the corresponding cell type. This conceptually simpler reformation in theory allows the model to generalize to any new TFs through their sequence and/or structural similarity with the TFs in the training data without the need of retraining it.

This reformulation also expands the original set of 1,903,668 labeled genomic bins into a substantially larger collection of individual *<*bin, TF, cell type*>* examples distributed across the training, validation, and test sets in Table 1. All 161 TFs and 91 cell types are represented in each split. As in TransBind, the new resulting dataset exhibits substantial class imbalance, with positive examples accounting for only 1.45%, 1.37%, and 1.36% of the training, validation, and test sets, respectively.

**Table 1.** Distribution of *<*DNA bin, transcription factor, cell type*>* examples used for training, validation, and testing in TransBind2.

| Split | Examples | % | % Positive |
| --- | --- | --- | --- |
| Train | 2,258,307,900 | 85.96 | 1.45 |
| Validation | 140,610,960 | 5.35 | 1.37 |
| Test | 228,142,980 | 8.68 | 1.36 |
| Total | 2,627,061,840 | 100.00 | – |

#### 1.1.2 Chromatin accessibility data

To provide information about chromatin accessibility in addition to the DNA sequence, TransBind2 uses DNase-seq data from ENCODE [11] uniformly processed DNase-seq bigWig files. For each 1,000 bp genomic bin, we extracted the per-base DNase signal from the bigWig track corresponding to the appropriate cell type, resulting in a 1,000-dimensional vector that is positionally aligned with the input DNA sequence. Positions without DNase-seq coverage were assigned a value of zero. DNase-seq data were available for 64 of the 91 cell types included in our dataset. Rather than excluding transcription factor–cell type pairs from the remaining 27 cell types without matching DNase-seq tracks, we assigned a fixed default signal value to preserve these samples during training. The resulting DNase signal was log-transformed and normalized prior to being provided as input to the model.

#### 1.1.3 Sequence uniqueness data

TransBind2 also incorporates genome uniqueness (mappability) information for each DNA bin, following the approach described in DeepGRN [6]. The Duke uniqueness track [11] assigns a score to each genomic position based on how uniquely the corresponding 35 bp sequence maps to the reference genome. Scores range from 0 to 1, where a value of 1 denotes a uniquely mappable sequence and lower values indicate decreasing uniqueness and increasing repetitiveness. For each 1,000 bp genomic bin, we computed the mean uniqueness score across the entire region to obtain a single scalar value. Unlike the DNase-seq signal, which preserves base-pair resolution, this scalar value was replicated across all 1,000 positions when constructing the model input.

#### 1.1.4 Transcription Factor Data

As in TransBind, TransBind2 uses the 161 unique transcription factors (TFs) present in the EPBDXDNA dataset, with amino acid sequences obtained from UniProt [30].

However, unlike TransBind, which uses ESM-DBP [36] to generate a single averaged embedding for each TF, TransBind2 employs ProstT5 [13], a protein language model designed to jointly capture sequence and structural information.

For each TF, a predicted three-dimensional structure was obtained from AlphaFold [16] and converted into a structural token sequence using Foldseek’s [31] 3Di alphabet. This represents the local structural environment of each residue as a discrete token. The amino acid sequence and corresponding 3Di token sequence were then independently encoded using ProstT5’s dual-prefix framework, producing 1,024-dimensional embeddings for each amino acid or 3Di alphabet at every residue position. These embeddings were concatenated to generate a combined 2,048-dimensional per-residue representation.

### 1.2 Deep Learning Model for TF-DNA Binding Site Classification

TransBind2 extends the TransBind architecture by incorporating chromatin accessibility (DNase-seq) and genome uniqueness (mappability) signals alongside the DNA sequence. It replaces the unidirectional cross-attention of TransBind with a symmetric, bidirectional cross-attention mechanism that lets DNA and protein representations mutually refine one another before prediction. The architecture retains a three-module decomposition: (i) DNA sequence encoder, (ii) protein encoder, and (iii) bimodal feature aggregation for TF–DNA binding prediction, as illustrated in Figure 1.

**Figure 1.**
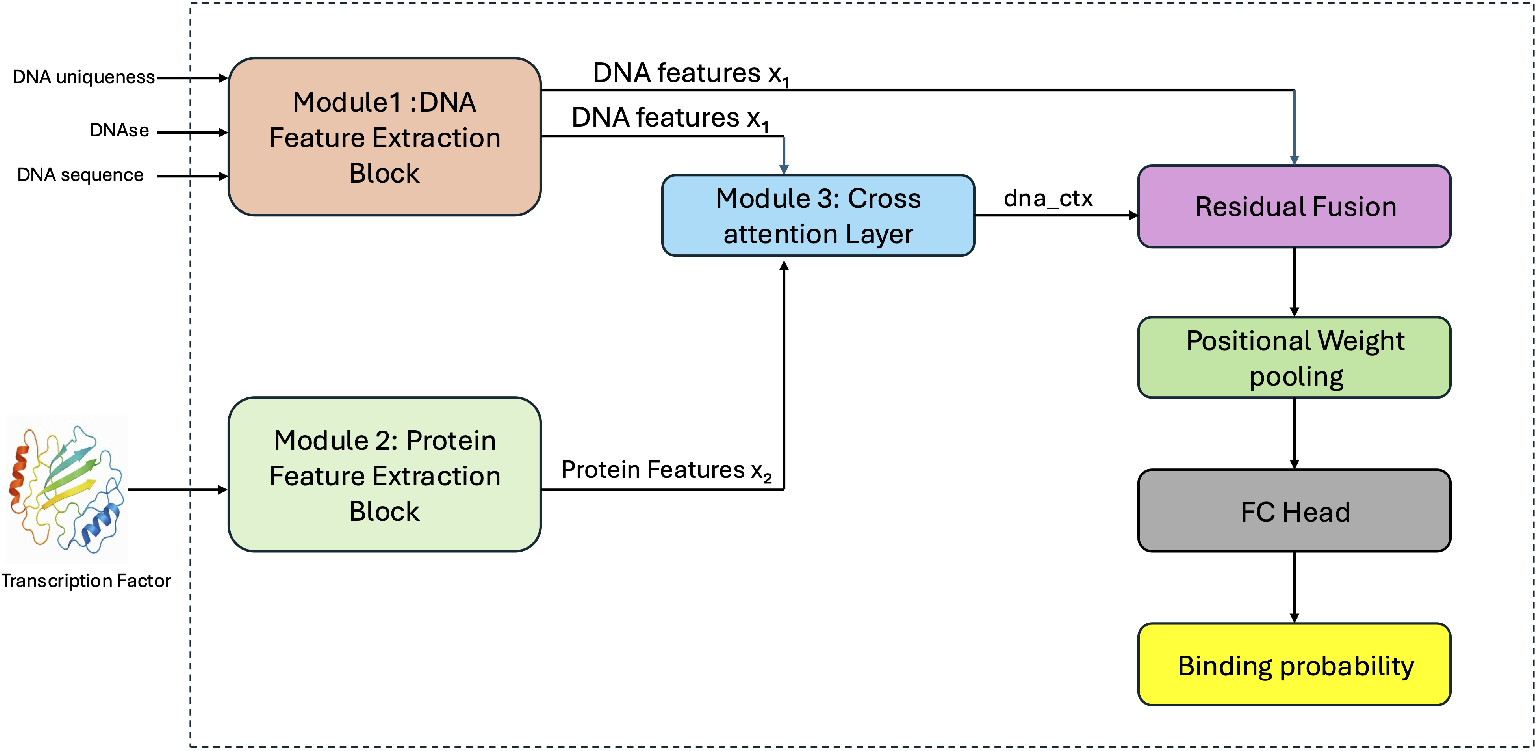
TransBind2 architecture for TF–DNA binding prediction. The model comprises three modules: (1) a six-channel DNA encoder that integrates sequence, DNase-seq accessibility, and mappability signals; (2) a protein encoder using ProstT5 embeddings with learned query cross-attention pooling; and (3) a bidirectional cross-attention fusion module that mutually refines DNA and protein representations before final TF-DNA prediction.

#### 1.2.1 Module 1: DNA Sequence Encoder

Each 1000-bp genomic bin is represented not only by its one-hot encoded DNA sequence but also by two auxiliary, position-aligned tracks: a DNase-seq accessibility signal and a genome uniqueness (mappability) signal as shown in Fig. 2. The one-hot encoded DNA sequence, consisting of four channels, is concatenated with the DNase and uniqueness tracks (one channel each) to form a six-channel DNA input tensor.

**Figure 2.**
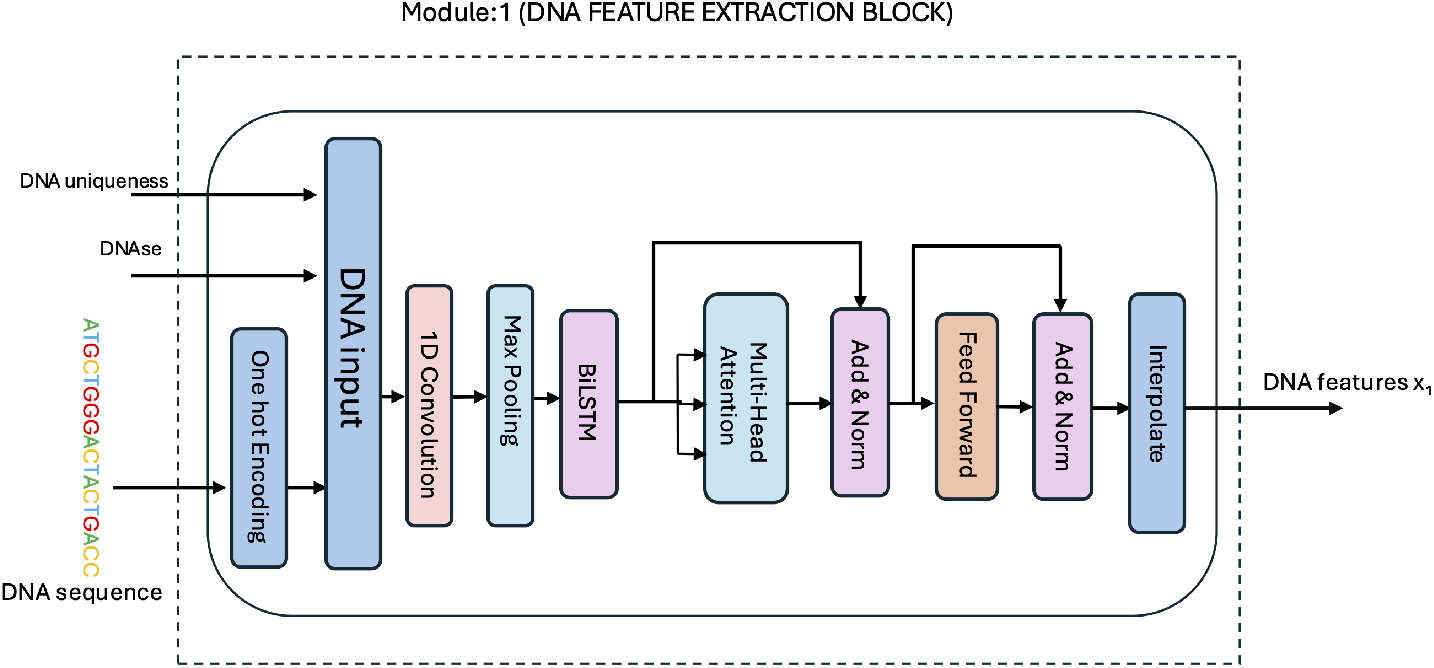
Module 1 (DNA sequence encoder): A 1000-bp genomic bin is represented as a six-channel tensor concatenating one-hot DNA encoding, DNase-seq accessibility, and genome mappability signals. Features are processed through convolutional and bidirectional LSTM layers, followed by a Transformer encoder with interpolation to a fixed 256-position representation.

The resulting six-channel tensor is processed by a one-dimensional convolutional layer (320 filters, kernel size 26), followed by ReLU activation and one-dimensional max pooling (window size 13, stride 13). The pooled features are then processed by a two-layer bidirectional long short-term memory (BiLSTM) network with 160 hidden units per direction.

The BiLSTM output is subsequently fed into a Transformer encoder layer consisting of 16 attention heads and a feed-forward network with a hidden dimension of 1024. The encoder follows the standard Transformer architecture, comprising a multi-head self-attention layer and a position-wise feed-forward network.

Unlike TransBind, which generates both a position-wise representation *H* and a separately pooled global embedding **x**_1_, TransBind2 does not compute an additional global summary vector. Instead, the position-wise output of the Transformer encoder is interpolated along the sequence dimension to a fixed length of 256 positions. The resulting tensor serves as the DNA feature representation (*x*_1_),

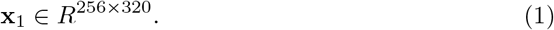

This representation is used as the DNA input to the cross-attention module in Module 3 and in the subsequent residual fusion step.

#### 1.2.2 Module 2: Protein Encoder

TransBind2 replaces the ESM–DBP [36] protein embedding used in TransBind with ProstT5 [13], a protein language model that jointly encodes amino acid sequence and structural information for each transcription factor (TF). For each TF, a predicted three-dimensional structure was obtained from AlphaFold [16] and converted into a 3Di structural token sequence using Foldseek [31]. ProstT5 then jointly processes the amino acid sequence and 3Di sequence, capturing both sequence-level and structural determinants of DNA-binding specificity as shown in Fig. 3. For each TF, ProstT5 generates a per-residue embedding with a dimensionality of 2048.

**Figure 3.**
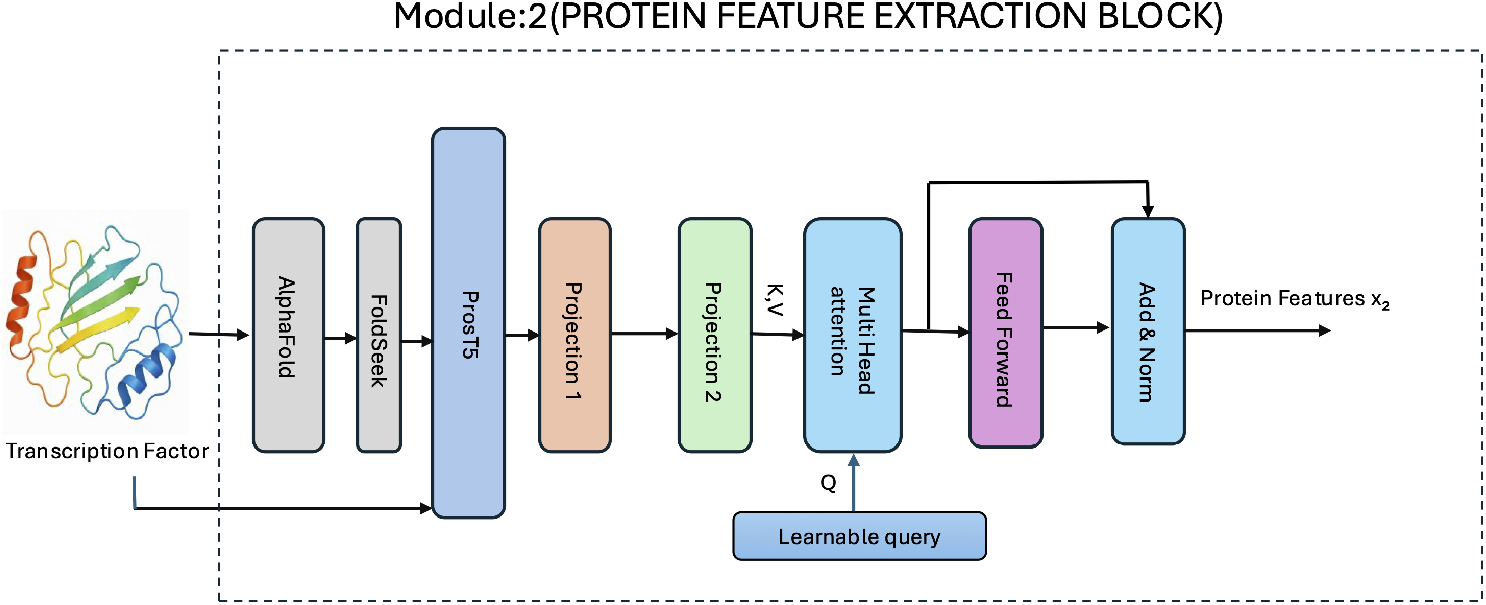
Module 2 (Protein encoder with learned query cross-attention pooling): ProstT5 jointly encodes amino acid sequence and AlphaFold-derived 3Di structural tokens for each TF, generating per-residue embeddings. A set of 160 learnable query vectors compress the variable-length protein embedding into a fixed 160×320 representation through scaled dot-product attention.

The variable-length embedding is compressed into a fixed-length representation using learned query cross-attention pooling. The embedding is first projected from 2048 to 512 dimensions and subsequently from 512 to 320 dimensions, with each projection followed by a Gaussian Error Linear Unit (GELU) activation function and dropout.

The resulting projected sequence serves as the keys and values for a multi-head attention operation, using the standard scaled dot-product attention mechanism,

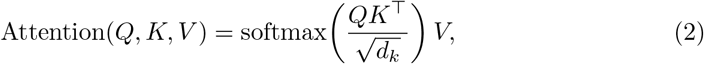

where *Q*, *K*, and *V* denote the query, key, and value matrices, respectively, and *d_k_*is the dimensionality of the key vectors. The same attention mechanism is used for the bidirectional cross-attention operations described in Module 3.

The queries are not derived from the protein sequence. Instead, they consist of a fixed set of 160 learnable query vectors that are shared across all TFs. These learnable queries define a fixed output length independent of the input sequence.

The attention output is processed by a position-wise feed-forward network followed by Add & Norm, where the residual connection is applied around the feed-forward sublayer. The resulting fixed-length protein representation (*x*_2_) is

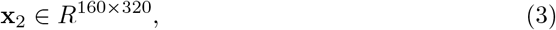

which consists of 160 protein token embeddings in the same 320-dimensional latent feature space as the DNA representation generated in Module 1, while remaining independent of the original TF sequence length.

#### 1.2.3 Module 3: Bimodal Feature Aggregation for TF–DNA Binding Prediction

As in TransBind, cross-modal interaction between DNA and protein representations is achieved through attention. However, as shown in Fig. 4, TransBind2 performs this interaction sequentially and bidirectionally rather than using a single one-directional attention mechanism.

**Figure 4.**
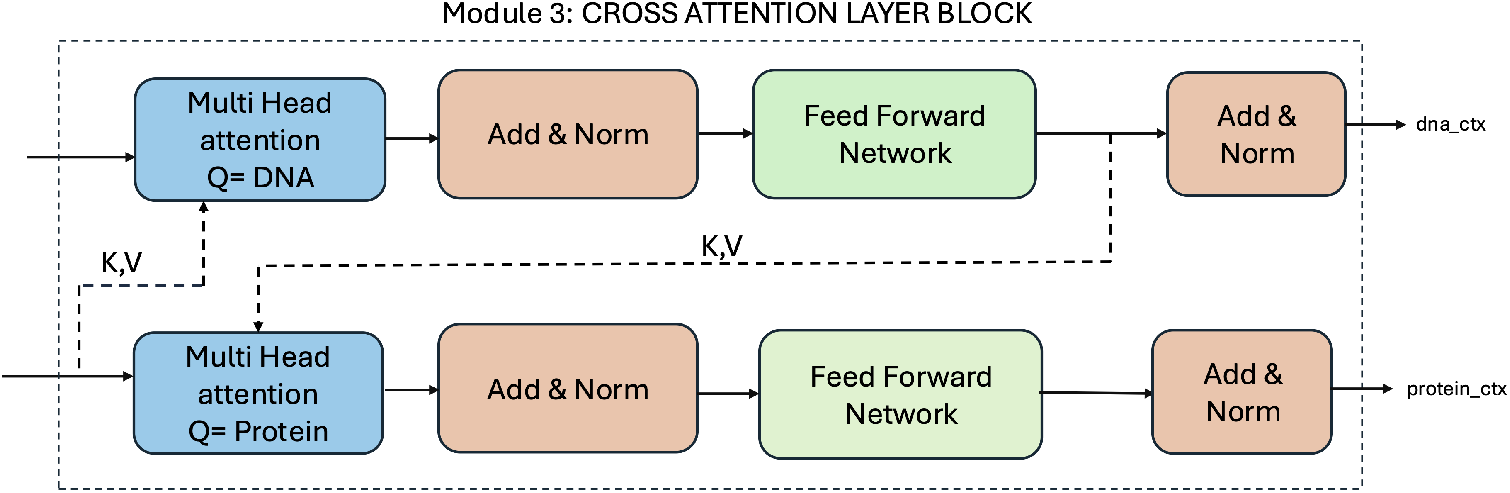
Module 3 (Bidirectional cross-attention fusion): DNA and protein representations are sequentially refined through bidirectional cross-attention. DNA features first attend to protein features to produce an updated DNA representation (**dna**_ctx_), which then serves as the key/value input for the protein-side update (**protein**_ctx_). This block is stacked for two successive layers.

The query, key, and value matrices are obtained through learned linear projections of their corresponding input representations. In the first operation, DNA features are used as queries and protein features as keys and values; the resulting attention output is followed by a residual Add & Norm connection, then passed through a position-wise feed-forward network with its own residual Add & Norm connection. This produces the updated DNA representation **dna**_ctx_. In the second operation, this direction is reversed: protein features are used as queries and **dna**_ctx_ as keys and values, followed by the same pattern of a residual Add & Norm connection after attention and another after the feed-forward network. This produces the updated protein representation **protein**_ctx_.

This sequential bidirectional update enables information exchange between the two modalities within each cross-attention layer. In the final architecture, this bidirectional cross-attention block is stacked for two successive layers, with both layers performing the protein-side update.

The final DNA representation, **dna**_ctx_, is then fused with the original DNA feature representation **x**_1_ from Module 1 through a residual fusion step (Figure 1). This residual connection preserves sequence-derived features while integrating information from the protein modality.

The fused DNA representation is subsequently processed by a positional weighted pooling layer, which learns a scalar attention score for each of the 256 sequence positions, normalizes these scores using a softmax function, and computes a weighted sum across positions to obtain a single pooled DNA representation. This adaptive pooling allows the model to emphasize sequence positions that are most informative for TF–DNA binding rather than pooling uniformly across the sequence.

Finally, the pooled representation is passed through a three-layer fully connected prediction head (320 → 512 → 256 → 1) followed by a sigmoid activation to produce the predicted TF–DNA binding probability.

### 1.3 Training and Evaluation

TransBind2 was implemented using PyTorch Lightning. It was trained on the training data and its hyperparameter optimization was performed on the validation data using Optuna (see Supplementary Note S1 for details). We additionally performed a hyperparameter importance analysis across all Optuna trials using fANOVA to assess the relative contribution of each hyperparameter to model performance (Supplementary Figure S1). The final model configuration used a learning rate of 3.20 × 10*^−^*^4^, a dropout rate of 0.057, weight decay of 0.034, a protein query length of 160 tokens, and two bidirectional cross-attention layers. The number of cross-attention heads was fixed at 8, with a model dimensionality of 320 across all optimization trials.

The final model was trained using the selected hyperparameter configuration. Because TF–DNA binding events are sparse relative to non-binding genomic regions, training batches were generated using a balanced sampling strategy with equal numbers of positive and negative examples in each batch (batch size = 3500). The model was optimized using binary cross-entropy with logits loss and the AdamW optimizer, with cosine annealing learning-rate scheduling and a minimum learning rate of 1 × 10*^−^*^6^.

Model selection was based on the checkpoint achieving the highest validation area under the precision-recall curve (AUPR), with early stopping applied after 15 consecutive epochs without improvement in validation AUPR.

Model performance was evaluated on the held-out test set (chromosomes 8 and 9) using the area under the receiver operating characteristic curve (AUROC) and the area under the precision-recall curve (AUPR). Predictions were compared with ground-truth labels separately for each TF experiment, and AUROC and AUPR values were then macro-averaged across TF experiments to obtain the overall reported performance, consistent with the evaluation procedure used for the other methods. AUROC measures ranking performance across classification thresholds and is unaffected by class prevalence, whereas AUPR was emphasized because of the strong class imbalance between TF–DNA binding and non-binding regions.

Finally, TrainsBind2 pretrained on the human data was blindly applied to a mouse dataset to assess how well it can generalize to new species and new TFs in a zero-shot prediction setting.

## 2 RESULTS

### 2.1 TransBind2 outperforms TransBind and existing state-of-the-art methods

We evaluated TransBind2 against the same set of baseline methods used in the original TransBind study, including DeepSEA [37], DanQ [23], TBiNet [22], finetuned DNABERT-2 [39], and EPBD×DNABERT-2 [17], together with TransBind [4] and TFBindFormer [18], a recently proposed attention-based model for TF–DNA binding prediction. All models were evaluated on the same held-out test set comprising chromosomes 8 and 9 using macro-averaged AUROC and AUPR across the 690 TF–cell type experiments. The EPBD×DNABERT-2 model could not be retrained due to unavailable training code and data; we therefore report its performance as described in the original publication.

As shown in Table 2, TransBind2 achieved the best performance among all methods, with an AUROC of 0.9648 and an AUPR of 0.4215. Relative to the second-best model, TransBind (AUROC 0.9508, AUPR 0.3741), this corresponds to relative improvements of 1.47% in AUROC and 12.67% in AUPR. The improvement in AUPR is particularly noteworthy given the severe class imbalance of TF–DNA binding labels (~1.44% positives), where AUPR provides a more informative and sensitive measure of performance than AUROC.

**Table 2.** AUROC and AUPR of TransBind2 and other methods on the test dataset. Bold font denotes the best result.

| Method | AUROC | AUPR |
| --- | --- | --- |
| DeepSEA | 0.8934 | 0.2509 |
| DanQ | 0.9254 | 0.3065 |
| TBiNet | 0.9402 | 0.3346 |
| finetuned DNABERT-2 | 0.9180 | 0.2960 |
| EPBD×DNABERT-2 | 0.9490 | 0.3260 |
| TFBindFormer | 0.9264 | 0.3291 |
| TransBind | 0.9508 | 0.3741 |
| **TransBind2** | **0.9648** | **0.4215** |

**Figure 5** presents the average ROC and PR curves across all 690 TF–cell type combinations for all methods. The ROC curves show that TransBind2 achieves higher true positive rates while maintaining lower false positive rates compared to all baseline methods. The PR curves indicate that TransBind2 obtains the highest precision across most of the recall range, with the largest separation from competing methods occurring at moderate-to-high recall, where baseline methods typically suffer most from false positives.

**Figure 5.**
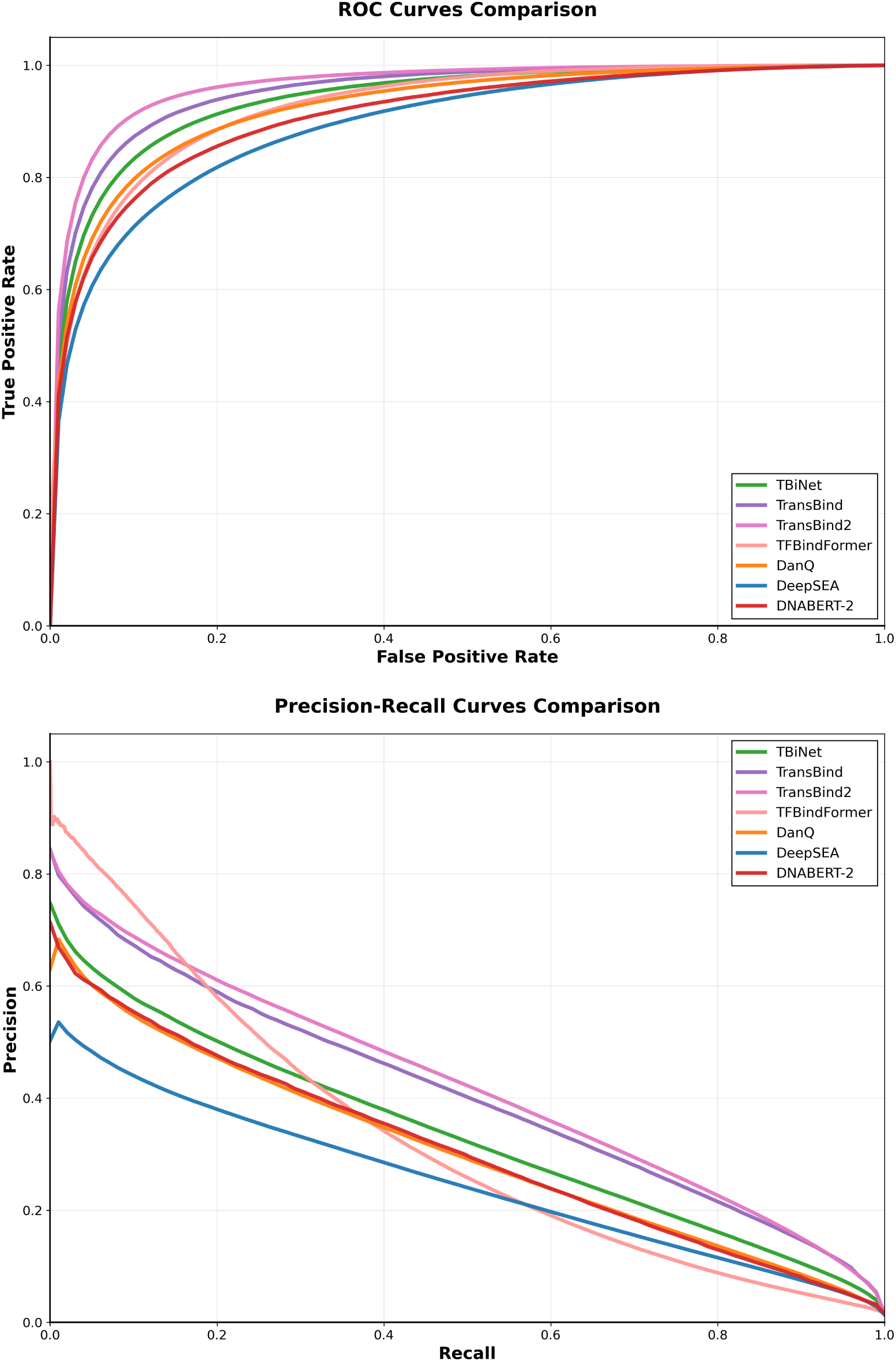
Average ROC curves (top) and Precision-Recall curves (bottom) across all TF-cell type combinations for all methods. TransBind2 demonstrates superior performance in both metrics, with particularly notable improvements in the precision-recall space.

To assess performance across individual TFs, we conducted a fine-grained comparison between TransBind2 and TransBind across all 161 unique TFs in the test dataset (Fig. 6), averaging scores across cell types to obtain a single AUROC and AUPR value per TF. TransBind2 outperformed TransBind in 156 of 161 TFs (96.9%) in terms of AUROC and 146 of 161 TFs (90.7%) in terms of AUPR, underscoring its robustness and broad applicability. Across all 161 TFs, TransBind2 achieved a mean AUROC of 0.96800.0207 compared to 0.9415 ± 0.0320 for TransBind, and a mean AUPR of 0.3631 ± 0.1445 compared to 0.2700 ± 0.1435 for TransBind. A paired *t*-test confirmed that these improvements are highly significant for both AUROC (mean difference 0.0265, *t* = 16.70, *p – value* = 6.71 × 10*^−^*^37^) and AUPR (mean difference 0.0931, *t* = 15.84, *p – value* = 1.35 × 10*^−^*^34^), with large effect sizes in both cases (Cohen’s *d* = 1.32 for AUROC and *d* = 1.25 for AUPR). A non-parametric Wilcoxon signed-rank test corroborated these results (AUROC *p – value* = 3.79 × 10*^−^*^27^; AUPR *p – value* = 1.39 × 10*^−^*^24^), confirming that the observed improvements are both statistically robust and practically meaningful. The individual performance curves for each TF-cell type experiment are provided in **Supplementary Figures S2 and S3**.

**Figure 6.**
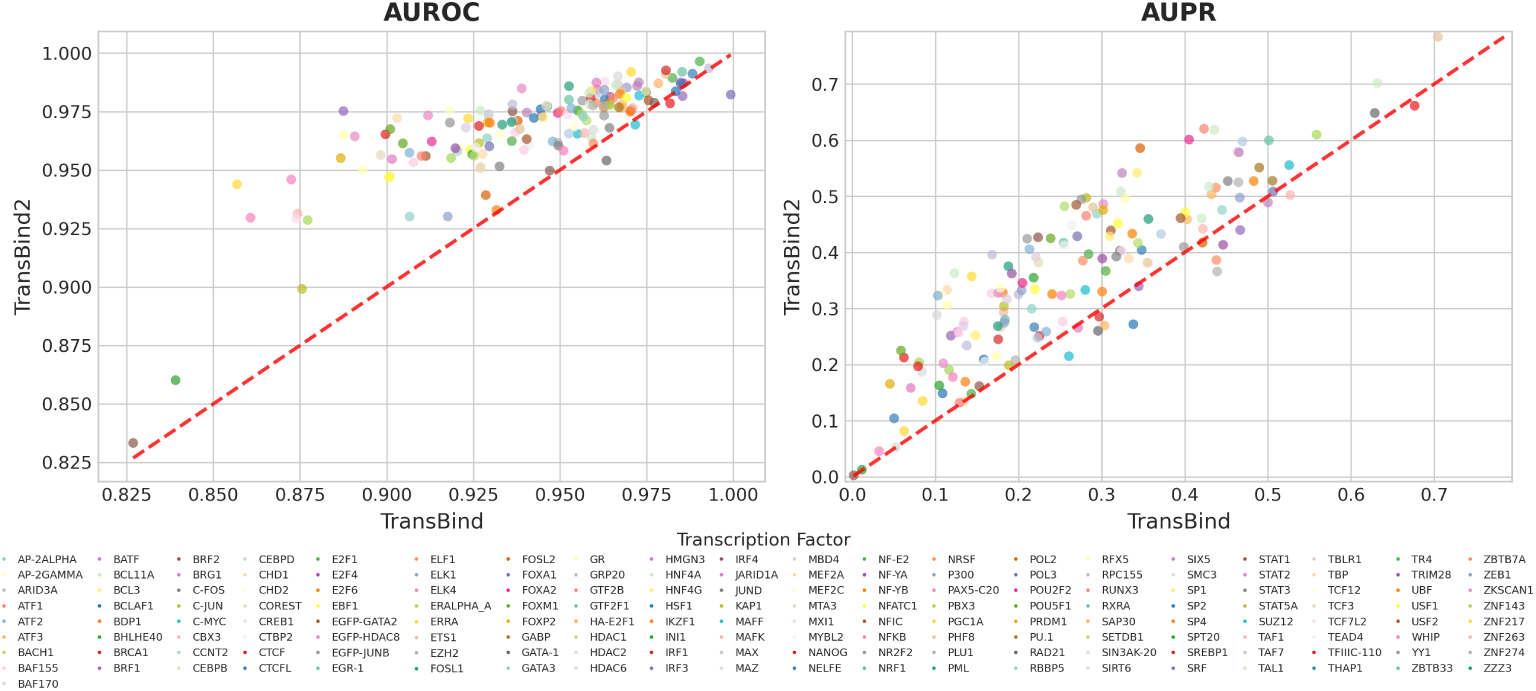
Performance comparison of TransBind2 versus TransBind across 161 TFs, aggregated from 690 TF–cell type experiments. Each point represents a TF in terms of AUROC (left) and AUPR (right). Points above the diagonal line indicate better performance by TransBind2. Each color denotes a unique TF.

The improvement is especially meaningful in biological contexts where the underlying data is highly imbalanced and the cost of false positives is high. For example, in many challenging cases where TransBind struggled, TransBind2 achieved dramatic gains, such as BCL11A (AUPR 0.123 → 0.363) and FOSL2 (AUPR 0.346 → 0.586). Even in many easier cases where TransBind’s performance was already good, TransBind2 still yielded notable improvements, e.g. CCNT2 (AUPR 0.469 → 0.598) and SMC3 (AUPR 0.632 → 0.702). Gains in AUROC followed a similar pattern, with the largest improvements observed for SRF (0.888 → 0.975), ZNF217 (0.857 → 0.944), and GR (0.888 → 0.975). Higher AUPR values translate into more precise identification of true binding sites, enabling more efficient downstream experimental validation.

### 2.2 TransBind2 generalizes across species: label-zero-shot prediction of mouse TF–DNA binding

The label-zero-shot evaluation in the original TransBind [4] study was limited to unseen TFs within the same species used for training (human). Here, we extended this evaluation to a cross-species setting by applying the human data-trained TransBind2 model directly to mouse ChIP-seq data [11]. To match the mouse system, we replaced species-specific inputs with mouse-derived features, including DNase-seq accessibility [11], CRG Alignability 36-mer mappability tracks [9, 11] on the mouse mm9 genome, and corresponding mouse TF structure representations [13, 16, 31].The DNA sequence input encoding strategy and the model architecture and weights were unchanged from the human data-trained TransBind2 model. This setting combines cross-species and label-zero-shot evaluation, as the model was applied to mouse genomic sequences and chromatin profiles not seen during training.

We evaluated TransBind2 on 26 TF–cell type combinations covering 22 unique TFs across two mouse cell lines (CH12 and MEL) (Table 3). Without access to mouse binding data during training, the model achieved a macro-averaged AUROC of 0.725 ± 0.088 and AUPR of 0.221 ± 0.071, substantially exceeding random prediction performance (AUROC = 0.5). Performance varied across TFs, likely reflecting differences in the conservation of TF–DNA binding mechanisms. The strongest AUROC values were observed for E2f4 (0.872), Sin3a (0.832 in CH12 and 0.816 in MEL), Hcfc1 (0.830), and Ubf (0.811). The highest AUPR values were obtained for Sin3a (0.398 in CH12), Gata1 (0.325 in MEL), and Mxi1 (0.305 in CH12).

**Table 3.** Cross-species label-zero-shot performance of TransBind2 on mouse TF–DNA binding prediction. Results include 26 TF–cell type combinations covering 22 unique TFs across two mouse cell lines (CH12 and MEL). AUROC, area under the receiver operating characteristic curve; AUPR, area under the precision–recall curve.

| TF | Cell type | AUROC | AUPR |
| --- | --- | --- | --- |
| Bhlhe40 | CH12 | 0.6571 | 0.2404 |
| Chd1 | CH12 | 0.7862 | 0.1226 |
| Chd2 | CH12 | 0.6447 | 0.1530 |
| Cmyc | CH12 | 0.6916 | 0.1381 |
| Corest | CH12 | 0.7661 | 0.1454 |
| Ctcf | MEL | 0.5799 | 0.1804 |
| Ctcf | CH12 | 0.5727 | 0.1943 |
| E2f4 | CH12 | 0.8720 | 0.0833 |
| Ets1 | CH12 | 0.7481 | 0.2863 |
| Gata1 | MEL | 0.7201 | 0.3251 |
| Hcfc1 | CH12 | 0.8300 | 0.2152 |
| Max | CH12 | 0.7131 | 0.2040 |
| Maz | CH12 | 0.7591 | 0.2476 |
| Mxi1 | CH12 | 0.7979 | 0.3050 |
| Nelfe | MEL | 0.7956 | 0.2094 |
| Nelfe | CH12 | 0.7793 | 0.2471 |
| P300 | CH12 | 0.7421 | 0.2418 |
| P300 | MEL | 0.7319 | 0.2094 |
| Pol2 | MEL | 0.6588 | 0.2991 |
| Pol2s2 | CH12 | 0.5233 | 0.2467 |
| Rad21 | CH12 | 0.6609 | 0.2276 |
| Sin3a | MEL | 0.8157 | 0.2730 |
| Sin3a | CH12 | 0.8324 | 0.3976 |
| Smc3 | CH12 | 0.7097 | 0.2444 |
| Tbp | MEL | 0.6459 | 0.1768 |
| Ubf | CH12 | 0.8110 | 0.1293 |
| **Overall** |  | **0.7248 ± 0.0876** | **0.2209 ± 0.0710** |

Overall, these results demonstrate that TransBind2 can generalize TF–DNA binding prediction across species boundaries, which has not been possible with previous deep learning methods in the field. The ability to predict mouse TF–DNA binding using a model trained entirely on human data suggests that the model captures conserved regulatory features across species.

### 2.3 Saliency maps localize TF–DNA binding signal within genomic windows

Although TransBind2 was trained using binary window-level binding labels, we asked whether the model learns where TF–DNA binding occurs within each 1000 bp genomic window (bin). To investigate this, we computed input × gradient saliency maps for correctly predicted bound examples in the test set and identified the nucleotide position with the highest saliency score within each window. We then compared these positions with experimentally determined ChIP-seq summit positions obtained from ENCODE [11] narrowPeak files, which provide high-resolution estimates of TF–DNA binding locations. When multiple ChIP-seq peaks (summits) overlapped a genomic window, the peak closest to the window center was used for evaluation.

We evaluated eight TFs spanning six structural families: CTCF (zinc finger, Cys2His2), EGR1 (zinc finger, Cys2His2), HNF4A (nuclear receptor), FOXA2 (forkhead), STAT3 (STAT), JUND (bZIP), YY1 (zinc finger, Cys2His2), and GATA3 (GATA zinc finger). For each TF, saliency-based predictions were compared against a center baseline that always predicts the window center (position 500 bp). Across all eight TFs, saliency maps substantially outperformed this baseline (Mann–Whitney *p <* 0.01 in all cases). Median positional errors ranged from 12–38 bp for saliency-based predictions (CTCF: 15 bp, EGR1: 16 bp, HNF4A: 22 bp, FOXA2: 17 bp, STAT3: 38 bp, JUND: 20 bp, YY1: 12 bp, GATA3: 34 bp), compared with 55–90 bp for the center baseline (CTCF: 66 bp, EGR1: 55 bp, HNF4A: 69 bp, FOXA2: 55 bp, STAT3: 90 bp, JUND: 75 bp, YY1: 74 bp, GATA3: 73 bp). Saliency-predicted positions were also strongly correlated with the corresponding ChIP-seq peak positions, with Pearson *r* = 0.699, 0.745, 0.784, 0.759, 0.755, 0.809, 0.698, and 0.616, respectively as shown in Fig. 7.

**Figure 7.**
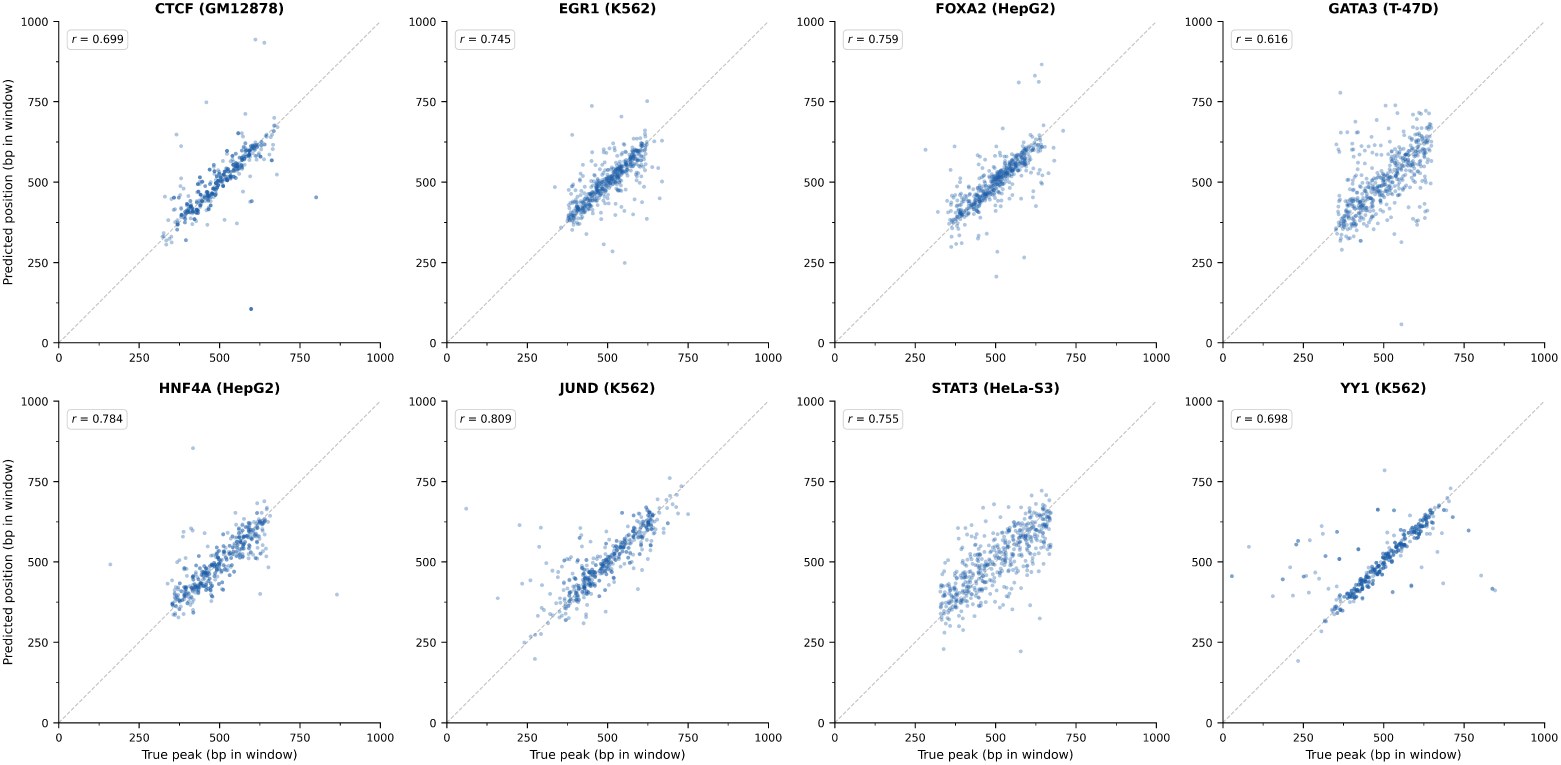
Saliency map localization of TF–DNA binding sites within genomic windows (bins). Each panel plots the position of peak localized by input × gradient saliency against the experimentally determined ChIP-seq peak position for 500 correctly classified TF-DNA binding examples in the test set. Results are shown for eight TFs spanning six structural families. The dashed line indicates perfect agreement between the saliency-localized peaks and the experimental peaks. Pearson correlation coefficients (*r*) are shown in each panel.

Despite receiving no position-level supervision during training, TransBind2 consistently identified positions close to the experimentally determined ChIP-seq peaks. The agreement between saliency-based predictions and ChIP-seq peak locations across eight TFs representing six structural families suggests that the model learns biologically meaningful positional information about TF–DNA binding events within genomic windows. In other words, the salience map-localized positions important for predicting binary TF-DNA binding events have a strong correlation with actual binding sites, indicating that the model implicitly learned the location of the binding sites to some extent even though no such labels were used in training.

## 3 Ablation Study of Input Modalities and Model Design

To quantify the contribution of each component of TransBind2, we performed two complementary sets of ablations. The first removes individual input modalities while holding the architecture fixed. The second holds the inputs fixed and varies how the protein and DNA representations are combined. Every ablated model was trained from scratch under identical settings and evaluated on the same held-out chromosomes (chr8 and chr9) across all 690 TF–cell type experiments. We report macro-averaged AUROC and AUPR. Because positive labels are extremely sparse, AUROC is compressed towards high values for all variants, and AUPR is the more discriminative measure.

### 3.0.1 Input modality ablations

We ablated each of the three information sources used by the full model: the ProstT5 protein embedding of the query transcription factor (**V1**), the DNase-seq accessibility channel (**V2**), and the mappability/uniqueness channel (**V3**). For V1, we removed the protein branch entirely; the model receives no TF information and predicts binding using only the DNA-derived representation, retaining the same binary prediction head as the full model. For V2 and V3, we removed the corresponding channel from the input tensor. This reduced the input from six to five channels, while keeping all other components unchanged. Results are summarised in Table 4.

**Table 4.** Effect of Input Modalities.

| Model variant | AUROC | AUPR |
| --- | --- | --- |
| TransBind2_V1 | 0.8843 | 0.1537 |
| TransBind2_V2 | 0.9430 | 0.3660 |
| TransBind2_V3 | 0.9630 | 0.4078 |
| **TransBind2 (final model)** | **0.9648** | **0.4215** |

The protein embedding is by far the most important input. Removing it (V1) reduces AUPR from 0.4215 to 0.1537, a relative loss of 63.5%. It is also the only ablation that causes a substantial drop in AUROC, from 0.9648 to 0.8843. This is consistent with the role of the protein branch in the model. Without a representation of TF, the model cannot distinguish between TFs and can only learn a general notion of regulatory potential shared across all TFs.

The DNase-seq channel has the second largest effect. Removing it (V2) reduces AUPR by 0.0555, or 13.2% relative to the full model. Accessibility provides information about chromosome states and cell types, which the protein embedding cannot capture. The same TF can bind to different genomic regions in different cellular contexts. A model that only uses sequence and protein identity has to average across these contexts.

The AUROC drop is comparatively small, from 0.9648 to 0.9430, a 2.3% relative decrease against the 13.2% relative decrease in AUPR. This suggests that accessibility mainly helps the model rank the strongest candidate sites more accurately.

The uniqueness channel has the smallest effect, but it still provides a consistent improvement. Removing it (V3) reduces AUPR from 0.4215 to 0.4078, a 3.3% relative drop. AUROC also decreases slightly, from 0.9648 to 0.9630.

Mappability helps the model identify regions where ChIP-seq read pile-ups may be unreliable. Its main benefit is therefore reducing false positives in repetitive and low-complexity regions. It has less effect on identifying additional true binding sites.

### 3.0.2 Multimodal fusion strategies

We tested how protein information is integrated with the DNA representation. All variants use the same six input channels and ProstT5 embeddings, but differ in how the TF and DNA modalities interact. In concatenation (**V4**), the protein representation is pooled into a single vector and concatenated with the DNA representation. In unidirectional cross-attention (**V5**), DNA positions attend to protein features, but not vice versa. The full model uses bidirectional cross-attention. Results are shown in Table 5.

**Table 5.** Effect of Multimodal Fusion Strategies.

| Model variant | AUROC | AUPR |
| --- | --- | --- |
| TransBind2_V4 | 0.9577 | 0.3791 |
| TransBind2_V5 | 0.9595 | 0.3906 |
| **TransBind2 (final model)** | **0.9648** | **0.4215** |

Performance improves as the interaction between protein and DNA becomes more expressive, going from V4 to v5 and to the final model. AUPR increases from 0.3791 with concatenation to 0.3906 with one-way attention and 0.4215 with bidirectional attention. Relative to the final model, this corresponds to AUPR drops of 10.1% for V4 and 7.3% for V5.

Because all variants use the same protein features, these differences reflect the fusion strategy itself. Moving from concatenation to one-way attention improves AUPR by 0.0115, while adding the reverse attention direction gives a further gain of 0.0309. The second step is therefore more than twice as large as the first, suggesting that allowing the protein representation to incorporate local DNA context is particularly important.

AUROC is much less affected, ranging from 0.9577 to 0.9648. The relative drops from the full model are only 0.7% for V4 and 0.5% for V5. Overall, the fusion strategy has a much larger effect on AUPR than AUROC, suggesting that richer protein–DNA interactions mainly improve the ranking of high-confidence binding sites rather than overall classification.

## 4 DISCUSSION

TransBind2 shows that modeling chromatin context and TF structure can improve genome-wide TF–DNA binding prediction. Bidirectional cross-modal attention further improves the model’s ability to combine these features. The 12.67% relative gain in AUPR over TransBind is especially important because AUPR is more informative than AUROC for highly imbalanced TF–DNA binding data. Some TFs, such as BCL11A and FOSL2, showed even larger gains. TransBind2 also improved AUROC or AUPR for most of the 161 TFs tested. This suggests that the new architecture provides a broad improvement rather than benefiting only a small subset of TFs.

The ablation results show how each component contributes to the model. The protein embedding has the largest impact. Removing it turns the model into a DNA-only encoder and causes the biggest drop in AUROC and AUPR. This shows that TF sequence and structural information are important for distinguishing binding from non-binding regions.

Chromatin accessibility has the second-largest effect. This is consistent with its role in capturing regulatory activity, chromosome states, and cell-type-specific binding. Its larger effect on AUPR than AUROC suggests that accessibility mainly helps the model identify true binding sites among many candidates.

Genome mappability has a smaller but consistent effect. It likely helps reduce false positives in repetitive or low-complexity regions rather than finding additional binding sites.

The fusion ablations also show that how the protein and DNA features are combined matters. Moving from simple concatenation to unidirectional attention, and then to bidirectional attention, leads to steady improvements in AUPR. Interestingly, adding the protein-to-DNA attention direction on top of the existing DNA-to-protein direction contributes more to performance than the DNA-to-protein direction does on its own, suggesting that allowing the protein representation to incorporate local DNA context is particularly valuable.

The cross-species zero-shot results show that TransBind2 can generalize beyond the human genome used for training. The model achieved macro-averaged performance clearly above random (AUROC = 0.725 vs. 0.5) on mouse ChIP-seq data. This suggests that it learns TF–DNA recognition patterns that are partly conserved across mammals, rather than simply memorizing human-specific sequence or chromatin patterns.

The saliency results provide another important finding. TransBind2 was trained using window-level binary labels, but its saliency maps can still identify the approximate location of TF-binding sites. The predicted positions consistently agree with experimentally measured ChIP-seq summits across TFs from different structural families. This suggests that the model learns meaningful information about binding-site location rather than treating each input window as a single unit. This could be useful for identifying regulatory elements and refining variant-effect predictions when base-pair-level experimental data are not available.

TransBind2 is trained and evaluated on 161 human TFs and 91 cell types. The pairwise reformulation of the prediction task provides a natural path for extending the model to additional TFs and cell types beyond this set. Because the model predicts binding for individual *<*DNA bin, TF, cell type*>* triplets rather than a fixed multilabel output, new ChIP-seq experimental data can be added to the training data directly, without retraining the model from a different architecture with a different number of outputs. Moreover, the cross-species test results on mouse ChIP-seq data already demonstrate this extensibility, and applying the same framework to additional species is the next step.

Building on these directions, future work could pursue three goals. First, joint training across multiple species could deepen this cross-species transfer, allowing the model to learn shared regulatory signals directly rather than relying on transfer from a model trained only on human data. Second, the framework could be extended to additional chromatin and epigenomic modalities broadening the biological context available to the model. Third, positional information such as ChIP-seq peak locations could be incorporated as an auxiliary training signal (labels), directly optimizing the model for base pair binding site resolution rather than relying on saliency maps computed after training.

## 5 DATA AVAILABILITY

TransBind2 is an open-source software package, and its source code is publicly available via Github at https://github.com/jianlin-cheng/TransBind2

## 6 ACKNOWLEDGEMENTS

This work is supported in part by funding from the National Science Foundation (NSF awards: # 2343612, # 2308699, and #2525780) and from the Department of Energy (award #: DE-SC0026121).

