## Supplementary Figures and Tables for "TransBind2: Improving Transcription Factor-DNA Binding Prediction with Multimodal Data and Bidirectional Cross Attention"

Supplementary Materials
Shreya Basnet[1,2], Jianlin Cheng[1,2,*]

**1** Department of Electrical Engineering & Computer Science, University of Missouri, Columbia, MO 65211, USA
**2** NextGen Precision Health, University of Missouri, Columbia, MO 65211, USA

### Hyperparameter Optimization and Training Configuration

#### Overview

We used the Optuna framework to tune the main architectural and training hyperparameters. Tree-structured Parzen Estimator (TPE) sampling was used to explore the search space, while MedianPruner was used to stop underperforming trials early. Hyperparameters were selected based primarily on validation AUPR, with AUROC used as a secondary evaluation metric.

#### Hyperparameter Search Space.

The following hyperparameters were included in the search:

- **Learning rate**: log-uniform distribution from $1.5 \times 10^{-4}$ to $5 \times 10^{-4}$
- **Dropout probability**: uniform distribution from 0.05 to 0.15
- **Weight decay**: log-uniform distribution from $1 \times 10^{-2}$ to $5 \times 10^{-2}$
- **Protein target length**: categorical choice of 96, 128, or 160
- **Number of cross-attention layers**: integer range from 2 to 3
- **Number of bidirectional layers**: integer range from 1 to the number of cross-attention layers

#### Selected Configuration.

The configuration with the best validation AUPR was selected:

- **Learning rate**: $3.20 \times 10^{-4}$
- **Weight decay**: 0.034
- **Dropout**: 0.057
- **Protein target length**: 160
- **Number of cross-attention layers**: 2
- **Number of bidirectional layers**: 2

This configuration was used for the final model and subsequent evaluation on the held-out test chromosomes.

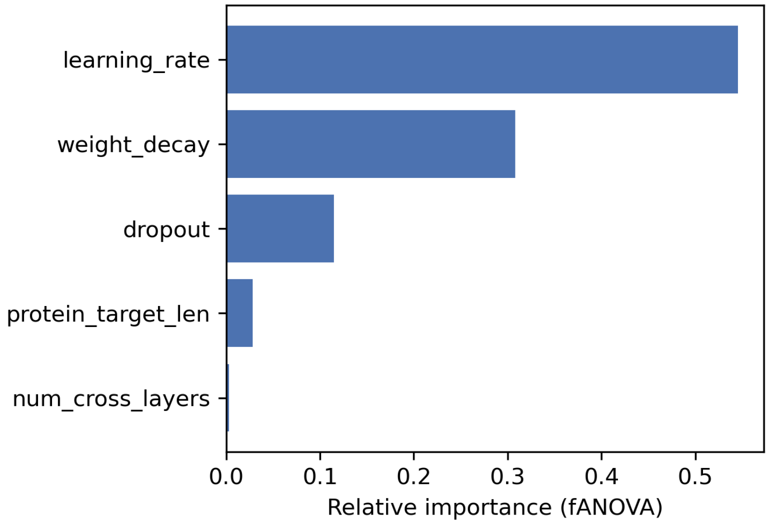


Figure S1: Hyperparameter importance analysis via fANOVA. Bars indicate the relative contribution of each hyperparameter to validation AUPR variance across completed Optuna trials. Learning rate and weight decay were the dominant drivers of performance, while architectural parameters had minor influence.

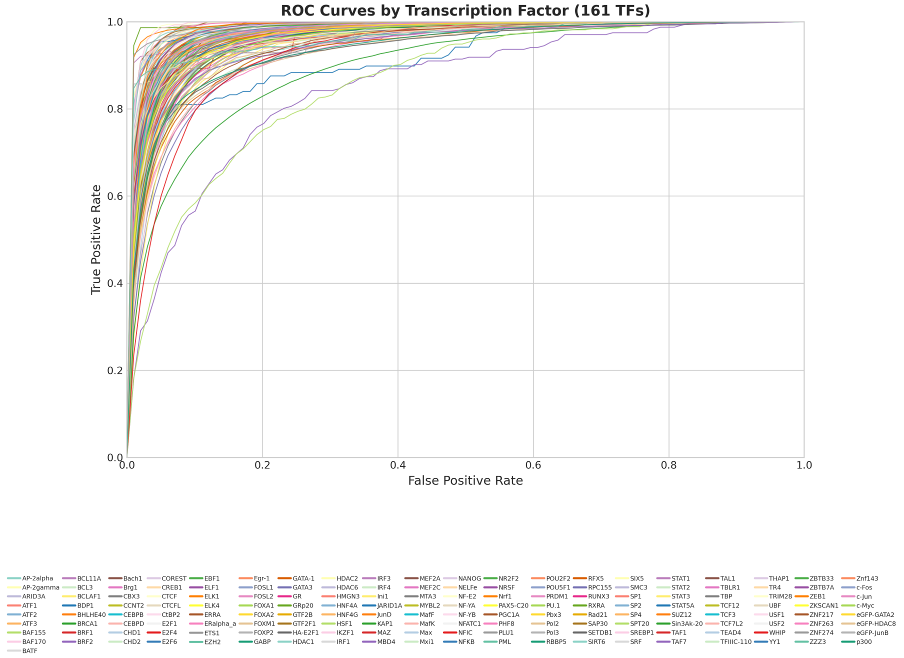


Figure S2: ROC curves for individual transcription factors in the test dataset. Each curve (colored distinctly) represents the performance for one unique TF, demonstrating the model's ability to discriminate DNA-binding sites across diverse transcription factor families.

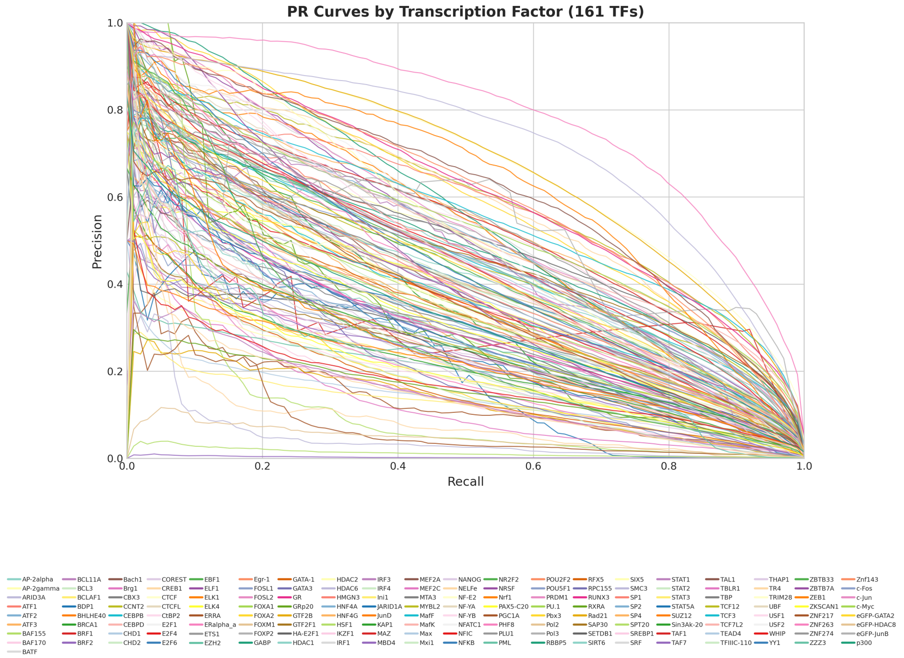


Figure S3: Precision-recall curves for individual transcription factors in the test dataset. Each curve (colored distinctly) represents one unique TF, illustrating the model's precision-recall trade-off across different transcription factor binding profiles.
